# A novel target associated with senescence and inflammatory signaling in human intervertebral disc degeneration

**DOI:** 10.64898/2026.09.02.748888

**Authors:** Máximo Alberto Diez-Ulloa, Iñaki López-Díaz, Adrián Cordido, Anxo Nogueira-Dabarca, José Ramón Caeiro-Rey, María D. Mayán

**Author notes:** Co-First Authors (Máximo Alberto Diez-Ulloa, Iñaki López-Díaz). **Corresponding author:** CellCOM Research Group. Centre for Research on Nanomaterials and Biomedicine (CINBIO). University of Vigo, and IISGS, SERGAS. Edificio Olimpia Valencia, Campus Universitario Lagoas Marcosende, 36310, Pontevedra, Spain., Complexo Hospitalario Universitario de Santiago de Compostela (CHUS). Universidade de Santiago de Compostela (USC). Santiago de Compostela, Spain.

## Abstract

**Background:** Intervertebral disc degeneration (IDD) is a leading cause of chronic low back pain and disability worldwide, affecting most individuals over 50 years of age. Despite its prevalence, no disease-modifying therapies exist, and current interventions are limited to reducing pain. Cellular senescence and the associated secretory phenotype (SASP) have been increasingly recognized as major drivers of disc matrix degradation and inflammation. However, the upstream molecular mechanisms that lead to IDD degeneration are still unknown. Connexin 43 (Cx43), a gap junction protein implicated in progression of age-related diseases, has emerged as a key regulator of cellular senescence and inflammatory signalling in musculoskeletal tissues.

**Methods:** Human primary cells were isolated from intervertebral disc samples obtained from patients classified into clinically meaningful groups: healthy controls, chronic/mechanical degeneration (DDD, ADJ, ASD), and acute/inflammatory event (herniated nucleus pulposus, HNP). Cx43 expression was assessed by qPCR and Western blotting. Cellular senescence was evaluated through SA-β-gal staining and analysis of p53/p21 expression. SASP factors and EMT-related markers were measured by qPCR. Protein expression was quantified by immunoblotting across different age groups and degeneration grades.

**Results:** In this current study Cx43, was identified as the most abundant connexin isoform in human intervertebral discs, showing a progressive increase in expression with age and disc degeneration. Also, high Cx43 expression correlated with increased expression of the senescent markers p53 and p21 and increased SA-β-gal activity. Besides, increased expression of EMT-related and differentiation markers has been correlated with high Cx43 levels in human IDD samples, consistent with fibrotic remodeling processes.

**Conclusions:** These findings identify aberrant upregulation of Cx43 signaling as a potential mechanistic link between intervertebral disc cellular senescence and extracellular matrix degradation, with the ensuing inflammatory response, representing a novel potential therapeutic target to modulate senescence-driven pathogenesis and modulate IDD progression.

## Introduction

Intervertebral disc degeneration (IDD) is a major contributor to chronic low back pain and spinal disability, being one of the most common musculoskeletal disorders affecting adults worldwide^1,2^. Epidemiological studies suggest that a high proportion of individuals over 50 show signs of disc degeneration, and many develop symptomatic disease with pain, reduced mobility, radiculopathy, and quality-of-life impairment^3^. Despite this burden, there is no approved disease-modifying therapy for IDD. Current interventions are limited to pain management, physiotherapy, or surgical procedures, none of which reliably restore disc structure or halt degeneration^4^.

Intervertebral disc degeneration (IDD) encompasses a spectrum of conditions that can be broadly divided into chronic, slowly progressive processes and acute mechanical disruptions. Degenerative Disc Disease (DDD) is characterized by progressive disc dehydration, loss of disc height, and annular fissuring^5^. Adjacent Level Disease (ADJ) refers to degenerative changes— including disc degeneration, stenosis, or instability—that develop at motion segments neighboring a fused spinal level^6^. Spondylolisthesis (SPL) involves vertebral slippage. There are two pathogenetic ways to SPL, one is developmental, occurs in growing children and adolescents and fall outside the scope of this research, the other is acquired, due to degenerative changes that concentrate in an specific level, usually L4-L5 due to biomechanical reasons: L5 is tightly connected to the pelvis so stresses concentrate at L4-L5. This acquired one, also referred to as degenerative spondylolisthesis, typically initiated by a monosegmental instability, which cause accelerated degeneration of the intervertebral disc or facet joints, leading to altered spinal balance and canal stenosis^7^. Adult Spinal Deformity (ASD, scoliosis) results from asymmetric disc and facet degeneration, causing a three-dimensional spinal curvature^8^; it is the results of multilevel segmental instability. In contrast, Herniated Nucleus Pulposus (HNP) represents an acute “blowout” of disc material beyond the annular boundary, triggering a rapid inflammatory response driven by cytokines such as IL-1β, TNF-α, and IL-6^9^. The previous chronic/mechanical degeneration also brings about an inflammatory response, but less explosive as it happens along several years or decades, is progressive and not triggered by a definite event, without a “date of beginning”, and as a difference, the inflammatory response is part of the process but not the triggering event (as in HNP)^9^.

IDD is a multifactorial, progressive process driven by mechanical, inflammatory, oxidative, and age-related stresses^10^. The intervertebral disc is composed of an inner nucleus pulposus (NP) and an outer annulus fibrosus (AF), with limited vasculature^11,12^. During the degeneration progress, disc height diminishes, the NP loses proteoglycans and water content, and the extracellular matrix (ECM) becomes stiffer and more degraded^13^ leading to pain and disability. Aberrant mechanical loading, microinjury, nutritional deficits, oxidative damage, and altered cell metabolism all contribute to disc degeneration and ECM breakdown^14^. Furthermore, cellular senescence is increasingly recognized as a convergent pathway in IDD^15^. Disc cells exposed to acidic pH, oxidative stress, nutrient deprivation, or chronic mechanical stress show increased markers of senescence, and adopt a senescence-associated secretory phenotype (SASP), releasing proinflammatory cytokines, matrix metalloproteinases (MMPs), and catabolic factors that further damage neighboring cells and ECM^16–18^. However, the upstream triggers and modulators of senescence in disc cells remain incompletely understood.

In recent years, connexins, particularly connexin 43 (Cx43), have been proposed to participate in the development of other musculoskeletal disorders such as osteoarthritis (OA)^19–23^. Connexins are a family of transmembrane proteins that assemble into hexameric hemichannels (connexons) on the plasma membrane. Paired hemichannels on adjacent cells form gap junctions (GJs), which allow direct intercellular exchange of small molecules (ions, second messengers, metabolites)^24,25^. In addition, hemichannels can mediate signaling to the extracellular space also contributing to tissue homeostasis. Connexins thus can regulate cell-to-cell coupling, paracrine signaling, and microenvironment homeostasis^26–28^. Our group has uncovered a novel therapeutic target, Cx43 involved in cellular senescence, inflammation and cartilage degradation in OA^19,22,23^. This transmembrane protein plays a pivotal role in promoting cellular senescence and chronic inflammation (inflammaging) in chondrocytes, synovial cells, and other joint tissues, driving OA progression^22^. Given this, we hypothesize that dysregulation of connexins, and particularly Cx43, in intervertebral disc cells may similarly play a role in IDD pathogenesis. Here, using human samples from IDD we have found that Cx43 expression progressively increases with age and intervertebral disc degeneration, in parallel with enhanced cellular senescence and SASP activity. This association suggests that dysregulated Cx43 signaling may play a key role in amplifying senescence and inflammatory processes within the intervertebral disc microenvironment.

## Results

To evaluate whether connexin expression is associated with intervertebral disc degeneration (IDD), human primary cells from patients were isolated from intervertebral disc samples and cultured in 2D conditions (Figure 1a). To account for the distinct pathobiological trajectories of intervertebral disc disease, samples were classified into three clinically and mechanistically meaningful groups. The Chronic/Mechanical group (DDD + ADJ + ASD) comprises slowly progressive degenerative processes, where endplate ossification compromises nutrient diffusion, creating a hypoxic, acidic niche that drives matrix remodeling and cellular senescence over months to years^29^. In contrast, the Acute/Inflammatory group (HNP) represents a sudden mechanical failure, a “blowout” of the nucleus pulposus through a ruptured annulus, that triggers an immediate innate immune response with elevated pro-inflammatory cytokines such as IL-1β, IL-6, and TNF-α^30^. This stratification allows us to dissect the molecular signatures (including connexin expression patterns) that distinguish slow, hypoxia-driven degeneration from the acute inflammatory cascade of herniation, a perspective that moves beyond the conventional focus on secondary inflammation in established disease.

**Figure 1.**
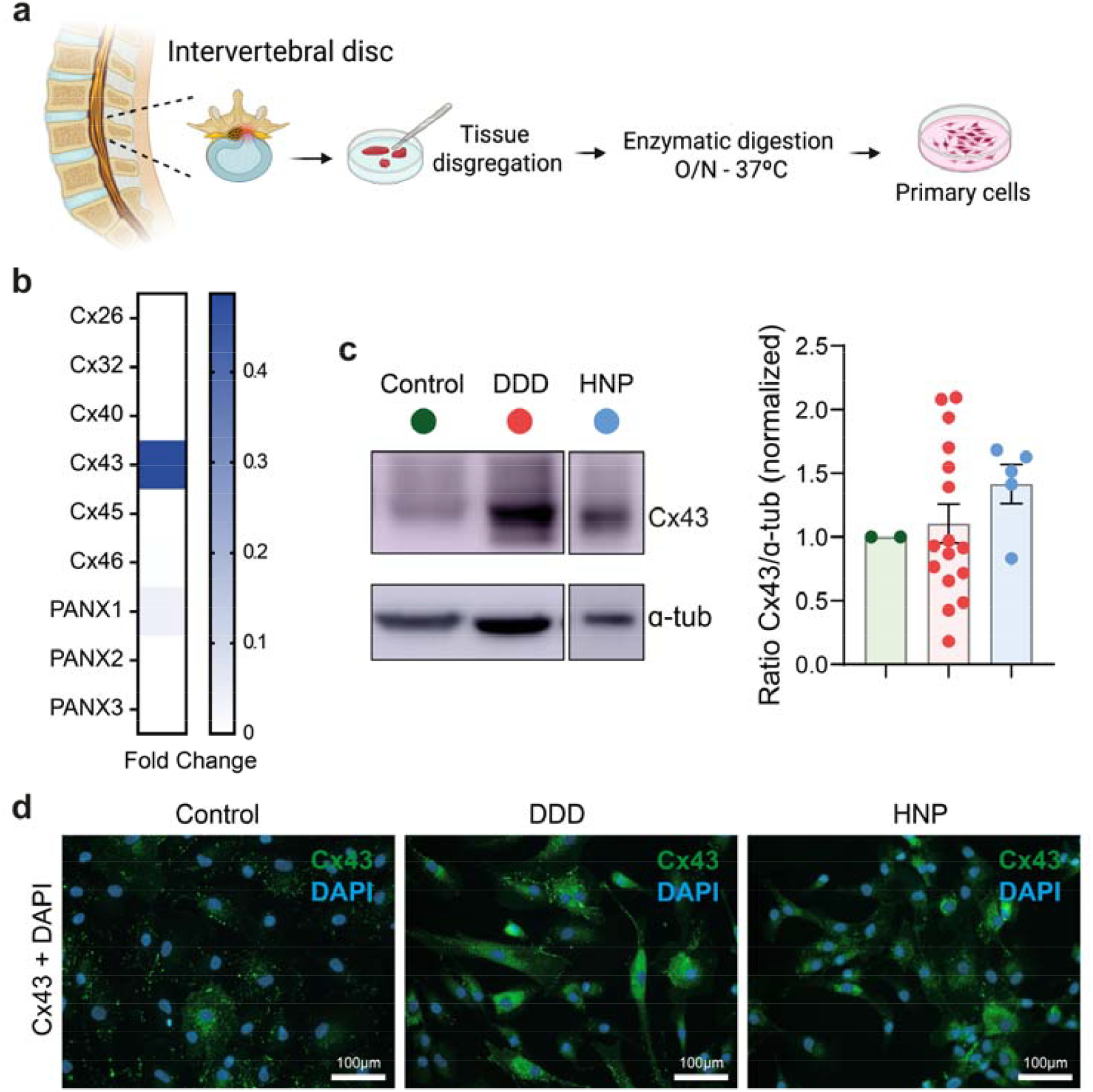
Expression of connexins and pannexins in the intervertebral disc. **a**. Diagram showing the steps to obtain disc primary cells isolated from human donors. **b**. RNA expression levels of different connexins (Cx) and pannexins (PANX) in vertebral cartilage human chondrocytes (vCH) derived from disc and endplate. Data were normalized to HPRT-1 levels (n = 3). **c**. Immunoblots from vCH lysates showing Cx43, levels in the indicated groups and its relative densitometric quantification. α-tubulin serves as loading control with mean of the Control group levels set to 1.0. Control means Healthy samples (n=2), DDD means Discal Degenerated Disc samples (n=14) and HNP means Herniated Nucleus Pulposus samples (n=5). d) Representative immunofluorescence for Cx43 staining in vCHs in the indicated groups (n=3 each). Scale bars 100μm. Statistical significance was determined by unpaired two-tailed Student’s t test, presented as mean ± s.e.m.

Firstly, we analyzed the expression of different connexin and pannexin genes in intervertebral disc tissue obtained from human donors classified as healthy (control) or degenerated (DDD and HNP). qPCR analysis of a panel of connexins and pannexins showed that, although Cx46, PANX1, and PANX2 were detectable in human disc tissue, Cx43 was the most abundantly expressed connexin and was consistently upregulated in degenerated discs, and the adjacent endplate region (Figure 1b). Cx43 protein levels positively correlated with mRNA levels showing upregulation in DDD and HNP groups compared with controls, suggesting that Cx43 upregulation may be associated with IDD phenotype (Figure 1c). Finally, we analyzed the localization and expression patter by immunofluorescence. The IDD samples of DDD and HNP groups showed a more intense Cx43 expression (Figure 1d).

We then analyzed Cx43 protein levels across different age ranges. Quantification of Western blot band intensities revealed a significant increase in Cx43 levels in the 40–51-year-old group compared with healthy samples, followed by a tendency toward decreased Cx43 levels in the 61–71-year age range (Figure 2a). In parallel, we examined cellular senescence across age groups in degenerated discs (DDD) by measuring senescence-associated β-galactosidase (SA-β-gal) activity. This analysis showed a progressive accumulation of senescent cells with donor ageing, increasing from discs from individuals aged 30–40 years to those older than 70 years, which displayed a significant increase in this senescence marker (Figure 2b). Notably, while Cx43 levels tended to not increase at advanced ages, SA-β-gal activity were significantly elevated, suggesting that senescence is maintained in aged discs (late adulthood) through mechanisms that may extend beyond Cx43 protein abundance. Next, we assessed the expression of the senescence-associated markers p53 and p21 at the protein level by Western blotting (Figure 3a). Both proteins showed a strong increased trend in degenerated discs compared with healthy samples.

**Figure 2.**
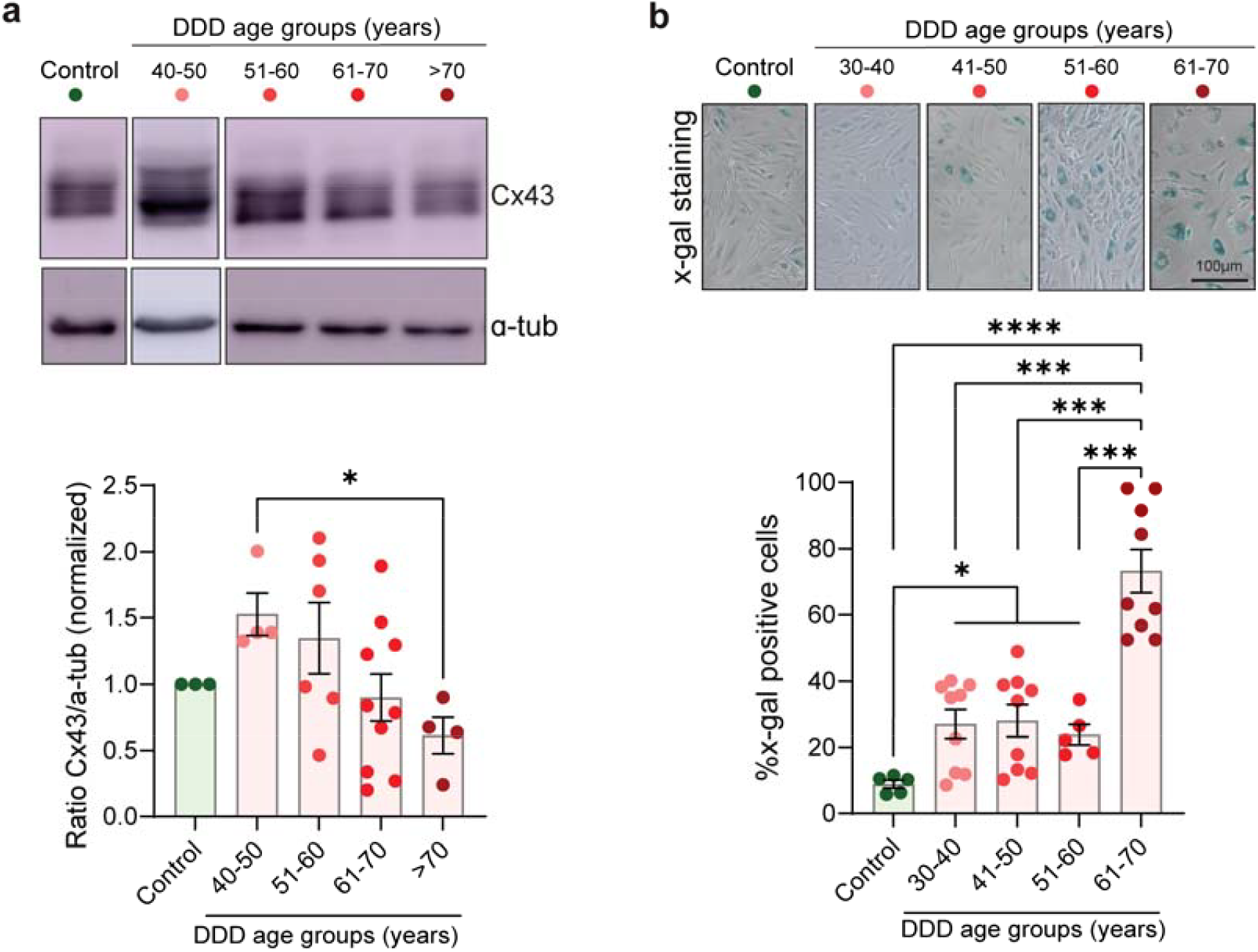
Correlation between Cx43 and senescence factors associated with tissue degeneration in this age-related disorder. **a**. Immunoblots from vCH lysates showing Cx43 protein from donor raging from healthy (Control) to Degenerative Disc Disease (DDD) patients in their in their 40-50 years (n=4), 51-60 years (n=6), 61-70 years (n=10) and >70 years (n = 4). α-tubulin serves as loading control with mean of the Control group levels set to 1.0. **b**. Senescence-associated β-galactosidase (β-gal) activity was analysed in vCHs derived from same group patients in their 30-40 years (n=9), 41-50 years (n=9), 51-60 years (n=5) and 61-70 years (n=9). On the upper pannel, representative images of the β-gal staining seen through a light. On the lower pannel, percentage of x-gal positive cells.Multiple-group comparisons are by one-way ANOVA followed by Brown-Forsythe and Welch tests multiple-comparison test, presented as mean ± s.e.m. *P <0.05, ***P<0.001, ****P<0.0001.

**Figure 3.**
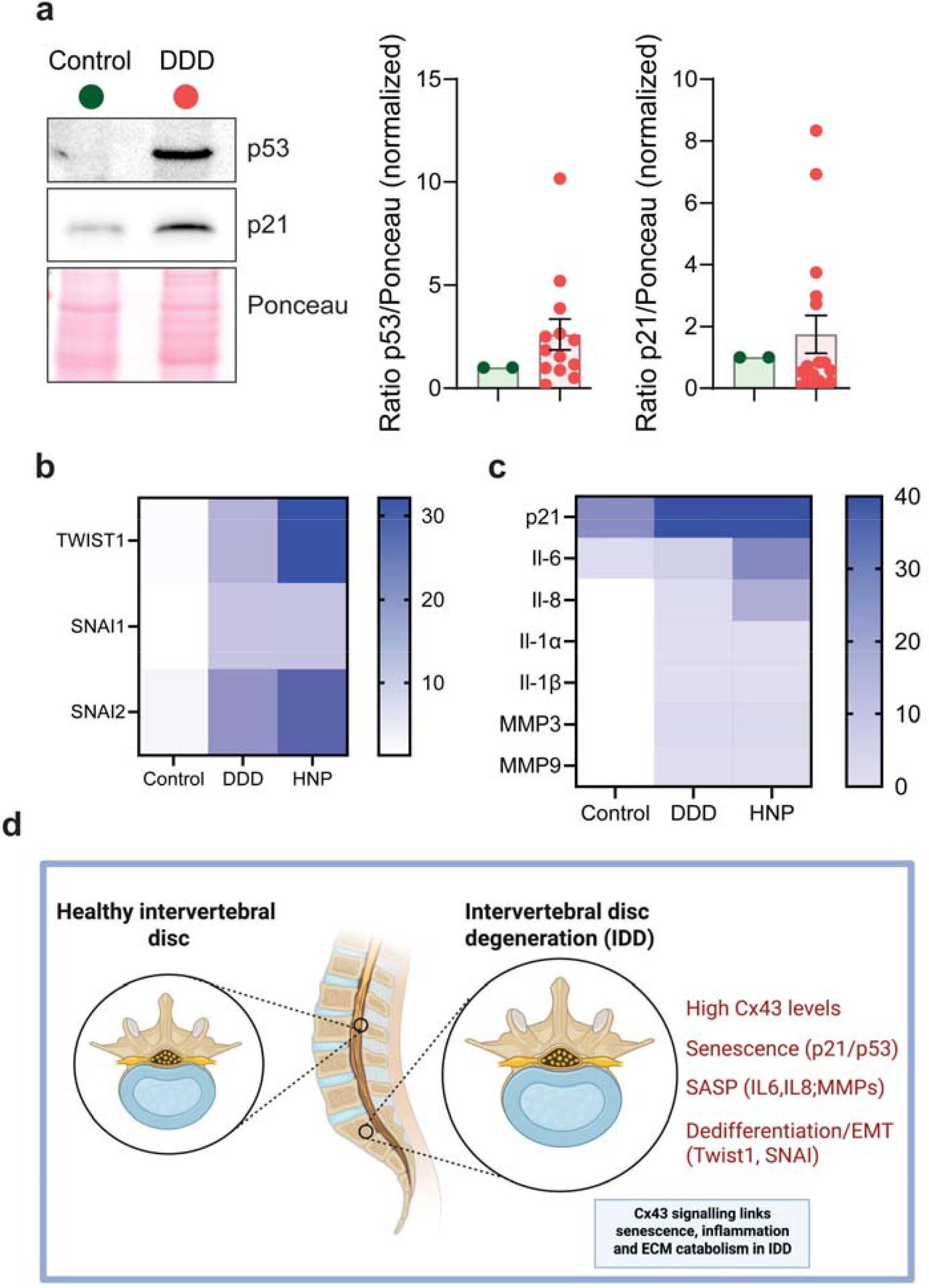
IDV shows a clear senescent phenotype. a) Immunoblots from vCH lysates, and its relative densitometric quantifications, showing p53 and p21 proteins from healthy (Control) to Degenerative Disc Disease (DDD) donors. Ponceau staining servers as loading control with mean of the Control group levels set to 1.0.**b, c**. Heatmap showing proinflammatory (*IL-6, IL-8, IL-1*α, *IL-1*β), matrix degradation (*MMP-3, MMP-9*), senescence (p21) and dedifferentiation (*TWIST1, SNAI1, SNAI2*) markers RNA expression levels analyzed in samples from healthy (Control), Degenerated Disc Disease (DDD) and Herniated Nucleus Pulposus samples (HNP) donors. White crosses represent the absence of expression. Data were normalized to HPRT-1 levels (n = ≥3; mean). **d**. Graphical abstract summarizing the results obtained. Cx43 is the most abundant connexin isoform expressed in human intervertebral disc tissue, and that its expression increases with age and degree of disc degeneration. Importantly, we showed that high levels of Cx43 corresponds with increased cellular senescence (SA-β-gal activity, p53/p21), enhanced expression of SASP factors (IL-6, IL-8, IL-1α/β, MMP-3, MMP-9) and up-regulation of dedifferentiation and EMT markers (TWIST1, SNAI1, SNAI2) in degenerated human discs. These results suggest that aberrant Cx43 signaling may play a mechanistic role in linking senescence, inflammation and ECM catabolism in IDD.

To further characterize the senescent and degenerative microenvironment, we performed qPCR analysis of senescence-associated secretory phenotype (SASP) factors. Degenerated discs showed increased expression of pro-inflammatory cytokines (IL-6, IL-8, IL-1α, IL-1β) and matrix-degrading enzymes (MMP-3 and MMP-9) (Figure 3b, c), consistent with a pro-inflammatory and catabolic tissue environment. In addition, degenerated samples exhibited increased expression of dedifferentiation and EMT-related transcription factors, including TWIST1, SNAI1, and SNAI2 (Figure 3b, c). These markers have been previously reported to be upregulated under conditions of enhanced Cx43 activity^22^, supporting the presence of phenotypic remodeling programs similar to those described in osteoarthritic chondrocytes^22^.

Collectively, these data demonstrate that Cx43 expression increases with disc degeneration and during mid-life aging, in parallel with cellular senescence, SASP activation, and loss of differentiated cell identity in IDD. At advanced ages, senescence persists despite the lack of increased Cx43 protein levels, indicating additional regulatory mechanisms contribute to disease progression in aged intervertebral discs.

## Discussion

In this study, we show that Cx43 (GJA1) is a predominant connexin detected in human intervertebral disc (IVD) tissue and that its expression associates with donor age and the degree of disc degeneration (IDD) samples. This observation is consistent with the fact that Cx43 is among the most ubiquitously expressed connexins in human tissues and is frequently implicated in age-related and degenerative pathologies^31^. In addition to Cx43, we detected Cx46 expression and members of the pannexin family, including *Panx1*, expanding the repertoire of channel-forming proteins that may contribute to disc homeostasis and degeneration. Notably, pannexins are increasingly recognized as regulators of inflammation and tissue remodeling in musculoskeletal contexts, and experimental evidence supports roles for pannexins (e.g., Panx3) in age-associated IVD degeneration^32^.

Our data indicate that high Cx43 levels correlate with multiple hallmarks of disc degeneration, particularly at ages associated with disease initiation (middle adulthood), including increased cellular senescence (SA-β-gal activity and p53/p21 activation), enhanced expression of key SASP factors (IL-6, IL-8, IL-1α/β), and increased expression of matrix-degrading enzymes (MMP-3, MMP-9). These findings align with work showing that senescent disc cells accumulate with age and degeneration and contribute to a pro-inflammatory, catabolic microenvironment that promotes extracellular matrix breakdown and disease progression^33^. We also observed upregulation of dedifferentiation and EMT-related transcription factors (TWIST1, SNAI1, SNAI2) in degenerated discs and involving in phenotypic remodeling in the context of degeneration-associated cell state transitions.

An additional point emerging from our age-stratified analysis is that, although Cx43 levels and SA-β-gal positivity track together in earlier age ranges, this relationship appears to weaken in older donors (e.g., ≥61 years). This divergence suggests that senescence may become sustained or amplified by Cx43-independent mechanisms at advanced ages, or that Cx43 function may be regulated at levels not captured by total protein abundance, including changes in subcellular localization, channel assembly/turnover, or hemichannel activity. These possibilities are consistent with broader literature emphasizing that connexin biology is strongly shaped by post-translational regulation and cellular context^34^.

The accumulation of senescent cells is well established in disc ageing and degeneration^16,33,35^. Senescent cells increase in number in aged and degenerated discs and likely drive degeneration through reduced proliferative capacity and chronic SASP release^16,36^. Our group has already linked Cx43 and senescence in other tissue contexts. Articular chondrocytes derived from OA cartilage contain high levels of Cx43 and this is associated with increased SASP, dedifferentiation and a pro□inflammatory environment^20–22^. Besides, we have demonstrated that extracellular vesicles enriched in Cx43 spread senescence, inflammation and reprogramming factors involved in wound healing failure to neighboring tissues, contributing to the progression of the disease among cartilage, synovium, and bone and probably from one joint to another^23^. Although cellular senescence in disc degeneration is well established as mentioned before and gap junctions in disc cells have been reported^37^ few studies until now have directly connected Cx43 up□regulation with senescence, SASP activation and dedifferentiation in human disc tissue. Also, previous findings are in line with earlier observations where it has been described that Cx43 (and Cx45) are expressed at the annulus fibrosus and participate in gap junction intercellular communication^37^. Thus, our data extend this knowledge by showing age and degeneration□associated up□regulation of Cx43 in human intervertebral disc tissue. Furthermore, in the musculoskeletal field, Cx43 has emerged as a key regulator of cell fate, mechanotransduction, aging and degenerative processes^19–22,38^. The present findings therefore place Cx43 into the IDD paradigm and raise the possibility that Cx43 is not only a marker of disc ageing but also may actively contribute to the degenerative cascade.

Overall, our results support a model in which aberrant Cx43 signaling is positioned at the intersection of senescence, inflammatory signaling, and ECM catabolism in IDD (Figure 3d). This framework is consistent with prior evidence linking Cx43 dysregulation to senescence-associated and inflammatory phenotypes in other degenerative contexts, including OA, and is compatible with emerging therapeutic concepts targeting Cx43-dependent pathways^25^. Taken together, these data encourage further investigation of Cx43 as a candidate therapeutic target to modulate degenerative signaling and potentially preserve disc cell function. Importantly, they also highlight the need to determine whether Cx43 exerts its effects through channel-dependent or channel-independent mechanisms, and to define the contribution of additional regulatory layers, including subcellular localization (e.g. nuclear versus membrane-associated channels), as previously suggested in recent studies^39^.

## Acknowledgements

This work was supported through funding 101079489-TWINFLAG and HORIZON-MSCA-2023-SE-01 to MDM funded by Horizon Europe, and grants PDI2022-137027OB-100 (MDM), funded by MICIU/AEI/10.13039/501100011033/ and by FEDER/EU, by GAIN, Xunta de Galicia 030_IN855A_2025-IGNICIA Programme. IL-D was funded by a predoctoral grant from Xunta de Galicia (IN606A-2022/036). Thanks to members of CellCOM and CINBIO for their support and helpful suggestions. To Marta Varela and Agustín Sánchez for their initial support.

## Author contributions

IL-D performed experiments, analyzed data, or prepared figures. AC analyzed data, prepared figures and contribute to manuscript revision. AND helped with manuscript revision and data analysis. MADU and JRCR provided critical input for conceiving the study, clinical advice and facilitated access to human samples. MDM directed and supervised the study. IL-D, AC and MDM wrote the manuscript with input from all co-authors. All authors reviewed and approved the final manuscript.

## Methods

### Primary cell extraction and cell culture

Vertebral human cartilage collection and processing were performed as previously described^20,21^. The study was conducted with the approval of the institutional Ethics Committee and after informed consent was obtained. Demographics, diagnosis, operated level(s), modified Pfirrmann grade of harvested IVDs are shown in Table S1 (Supportive information). We selected vertebral cartilage samples from patients with moderate and advanced IDD from both sexes.

We stratified our IDD samples into three groups: (A) Healthy controls (Control); (B) Chronic/Mechanical degeneration (DDD + ADJ + ASD), encompassing established, slowly progressive degenerative processes driven by matrix remodelling, and mechanical overload^40^; and (C) Acute/Inflammatory HNP (HNP), representing an acute biomechanical “blowout” with annular rupture and a pronounced early inflammatory response^30^. This classification acknowledges that while DDD and HNP share some biochemical features, they represent distinct temporal and biomechanical entities.

Cartilage human chondrocytes (vCH) derived from IDD patient discs, were isolated from fresh tissue as previously described^21^. Cells (2×10^6) were seeded into 100-mm dishes and incubated at 37 °C with 5% CO2 and 100% humidity in Dulbecco’s Modified Eagle’s Medium (DMEM) supplemented with 100 U/ml penicillin, 100 μg/ml streptomycin, and 10% foetal bovine serum (FBS; all from Gibco, Thermo Fisher Scientific) until ~80–90% confluence was reached.

### Immunofluorescence assay

Cells were seeded onto coverslips and incubated overnight to allow for attachment. Following treatment, the culture medium was removed and the cells were washed with PBS. Cells were fixed with 4% paraformaldehyde (PFA; Sigma-Aldrich, #158127-100G) in PBS at room temperature (RT) for 10 minutes. The fixative was removed, and the samples were washed again with PBS. To quench autofluorescence, 0.1 M glycine (Sigma-Aldrich, #G7126-1KG) in PBS was added to each well for 15 minutes at RT. Membrane permeabilization was performed using PBS containing 0.25% Triton X-100 (Sigma-Aldrich, #93443-100ML) for 10 minutes at RT. Non-specific binding sites were blocked with a solution of 1% bovine serum albumin (BSA; Sigma-Aldrich, #A9418-50G) and 0.1% Tween-20 (Sigma-Aldrich, #P2287-500ML) in PBS (PBST) for at least 30 minutes at RT. Samples were incubated with the Cx43 primary antibody (1:200, Sigma-Aldrich, #C6219) overnight at 4°C in a humidified chamber. After washing with PBS, cells were incubated with goat anti-rabbit Alexa Fluor 594 secondary antibody (1:200, Thermo Scientific #A32740) for 1 hour at RT in the dark. Following further PBS washes, nuclei were counterstained with DAPI (1:1000 in PBS) for 5 minutes at RT. After a final wash, coverslips were mounted using Glycergel mounting medium and visualized in a Nikon Eclipse Ti fluorescence microscope.

### Protein extraction and immunoblot analysis

Cells were lysed on ice using lysis buffer (150 mM NaCl, 50 mM Tris, pH 8, 50 mM EDTA, pH 8, 0.5% v/v Nonidet P-40, 0.3% v/v sarkosyl, and 0.1% w/v SDS), supplemented with 5 μg/ml protease inhibitor cocktail and 1 mM phenylmethylsulfonyl fluoride (PMSF; Sigma-Aldrich, #P7626). Protein concentration was determined with the Pierce™ BCA Protein Assay Kit (Thermo Fisher, #23225). For electrophoresis, 20 μg of total protein were mixed with loading buffer at a 5:1 (v/v) ratio (10% SDS, 200 mM Tris-HCl pH 6.8, 50% glycerol, 0.1% bromophenol blue, and 10% β-mercaptoethanol) and separated on a 10% SDS-PAGE gel. Proteins were transferred to a polyvinylidene fluoride (PVDF) membrane (Millipore Co., #IPVH00010). Membranes were first stained with ATX Ponceau S red staining solution (Sigma-Aldrich, #09189-6X1L-F), then blocked in 5% milk prepared in PBS with 0.05% Tween-20 (Sigma-Aldrich, #P2287). After blocking, membranes were incubated overnight at 4 °C with primary antibodies, washed, and further incubated with an HRP-conjugated secondary antibody at room temperature for 1 h. Detection was carried out using the Pierce™ ECL Western Blotting Substrate and visualized in an Amersham Imager 600. The following primary antibodies were used: α-tubulin (1:1000, Sigma-Aldrich, #T9026),Cx43 (1:1000, Sigma-Aldrich, #C6219), p53 (1:500, Santa Cruz Biotechnology, #sc-126) and p21 (1:500, Cell Signaling Technology, #2947S).

### Real-time quantitative qPCR gene expression

Total RNA was isolated from cells using TRIzol™ reagent (Invitrogen, Thermo Fisher Scientific, #15596018) following the manufacturer’s protocol. One microgram of total RNA was used for cDNA synthesis with the SuperScript® VILO™ cDNA Synthesis Kit (Invitrogen, Thermo Fisher Scientific, #12013679). Quantitative PCR was performed using the Applied Biosystems™ PowerUp™ SYBR™ Green Master Mix (Thermo Fisher Scientific) on a real-time PCR system (LightCycler® 480, Roche) with the primers listed in Table 2.

**Table 2.**
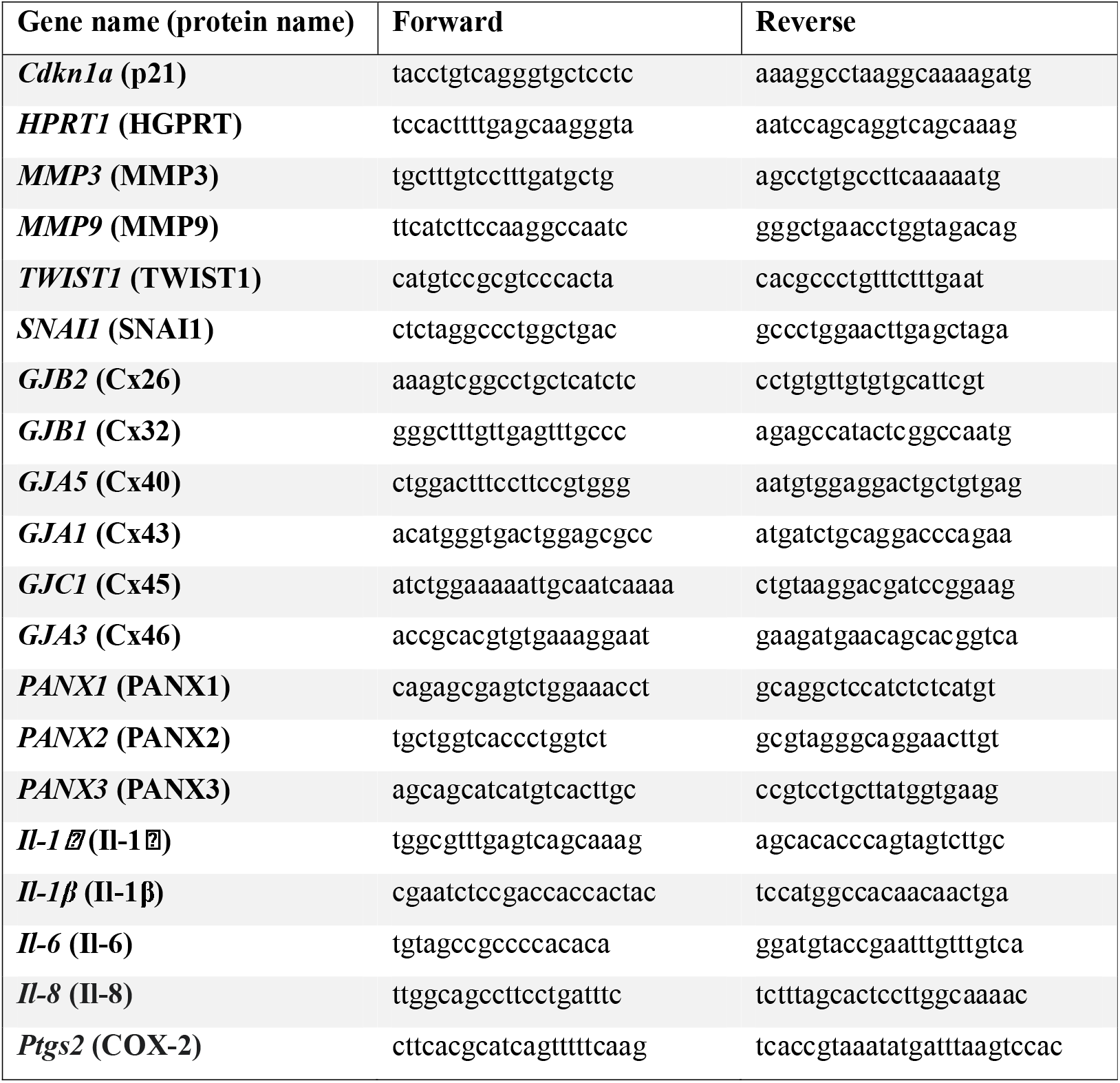
RT-PCR primer sequences.

### Senescence-associated β-galactosidase staining

Cells were seeded in a 12-well plate left for 72 hours. SA-β-Gal activity was analysed using the Senescence Cells Histochemical Staining Kit (Sigma-Aldrich, #CS0030). Briefly, cells were rinsed with warm PBS, fixed, and incubated overnight (O/N) at 37 °C without CO□in a staining solution containing X-gal. Cells were imaged with a Nikon Eclipse TS100 microscope, and the percentage of SA-β-Gal–positive (blue) cells was quantified using ImageJ software (version 1.53k).

### Statistics

All statistical analyses were performed using GraphPad Prism (version 8.4.). Data are presented as mean ± s.e.m. Differences between the two groups were assessed using unpaired two-tailed Student’s t-tests. Comparisons among multiple groups were performed using one-way analysis of variance (ANOVA) with Brown-Forsythe and Welch post hoc test. Statistical significance was set at P < 0.05 (*P<0.05, **P<0.01, ***P<0.001).

## References

1. Samanta A, Lufkin T, Kraus P. Intervertebral disc degeneration-Current therapeutic options and challenges. Front Public Health. 2023;11:1156749. doi:10.3389/fpubh.2023.1156749

2. Kirnaz S, Capadona C, Wong T, et al. Fundamentals of Intervertebral Disc Degeneration. World Neurosurg. Jan 2022;157:264–273. doi:10.1016/j.wneu.2021.09.066

3. Zhang X, Zhang Z, Zou X, et al. Unraveling the mechanisms of intervertebral disc degeneration: an exploration of the p38 MAPK signaling pathway. Front Cell Dev Biol. 2023;11:1324561. doi:10.3389/fcell.2023.1324561

4. Xin J, Wang Y, Zheng Z, Wang S, Na S, Zhang S. Treatment of Intervertebral Disc Degeneration. Orthop Surg. Jul 2022;14(7):1271–1280. doi:10.1111/os.13254

5. Gradisnik L, Kocivnik N, Maver U, Velnar T. Degenerative Disease of Intervertebral Disc: A Narrative Review of Pathogenesis, Clinical Implications and Therapies. Bioengineering (Basel). Dec 29 2025;13(1)doi:10.3390/bioengineering13010040

6. Helgeson MD, Bevevino AJ, Hilibrand AS. Update on the evidence for adjacent segment degeneration and disease. Spine J. Mar 2013;13(3):342–51. doi:10.1016/j.spinee.2012.12.009

7. Saremi A, Goyal KK, Benzel EC, Orr RD. Evolution of lumbar degenerative spondylolisthesis with key radiographic features. Spine J. Jun 2024;24(6):989–1000. doi:10.1016/j.spinee.2024.01.001

8. Lee JH, Lee SH, Cho EA, et al. Asymmetric Degenerative Changes Between Convex and Concave Sides in Symptomatic Adult Degenerative Scoliosis. J Clin Med. Sep 26 2025;14(19)doi:10.3390/jcm14196825

9. Yussen PS, Swartz JD. The acute lumbar disc herniation: imaging diagnosis. Semin Ultrasound CT MR. Dec 1993;14(6):389–98. doi:10.1016/s0887-2171(05)80032-0

10. Hammoor BT, Lai CS, Xiong GX, et al. Intervertebral disc degeneration. Nat Rev Dis Primers. Feb 5 2026;12(1):5. doi:10.1038/s41572-025-00681-8

11. Podichetty VK. The aging spine: the role of inflammatory mediators in intervertebral disc degeneration. Cell Mol Biol (Noisy-le-grand). May 30 2007;53(5):4–18.

12. Yurube T, Takeoka Y, Kanda Y, Kuroda R, Kakutani K. Intervertebral disc cell fate during aging and degeneration: apoptosis, senescence, and autophagy. N Am Spine Soc J. Jun 2023;14:100210. doi:10.1016/j.xnsj.2023.100210

13. Ke W, Xu H, Zhang C, et al. An overview of mechanical microenvironment and mechanotransduction in intervertebral disc degeneration. Exp Mol Med. Oct 01 2025;doi:10.1038/s12276-025-01546-6

14. Liu F, Chao S, Yang L, et al. Molecular mechanism of mechanical pressure induced changes in the microenvironment of intervertebral disc degeneration. Inflamm Res. Dec 2024;73(12):2153–2164. doi:10.1007/s00011-024-01954-w

15. Aasen T, Leithe E, Graham SV, et al. Connexins in cancer: bridging the gap to the clinic. Oncogene. 06 2019;38(23):4429–4451. doi:10.1038/s41388-019-0741-6

16. Silwal P, Nguyen-Thai AM, Mohammad HA, et al. Cellular Senescence in Intervertebral Disc Aging and Degeneration: Molecular Mechanisms and Potential Therapeutic Opportunities. Biomolecules. Apr 18 2023;13(4)doi:10.3390/biom13040686

17. Chen X, Zhang A, Zhao K, et al. The role of oxidative stress in intervertebral disc degeneration: Mechanisms and therapeutic implications. Ageing Res Rev. Jul 2024;98:102323. doi:10.1016/j.arr.2024.102323

18. Song C, Zhou Y, Cheng K, et al. Cellular senescence - Molecular mechanisms of intervertebral disc degeneration from an immune perspective. Biomed Pharmacother. Jun 2023;162:114711. doi:10.1016/j.biopha.2023.114711

19. Carpintero-Fernández P, Varela-Eirín M, García-Yuste A, López-Díaz I, Caeiro JR, Mayán MD. Osteoarthritis: Mechanistic Insights, Senescence, and Novel Therapeutic Opportunities. Bioelectricity. Mar 2022;4(1):39–47. doi:10.1089/bioe.2021.0039

20. Mayan MD, Carpintero-Fernandez P, Gago-Fuentes R, et al. Human articular chondrocytes express multiple gap junction proteins: differential expression of connexins in A normal and osteoarthritic cartilage. Am J Pathol. Apr 2013;182(4):1337–46. doi:10.1016/j.ajpath.2012.12.018

21. Mayan MD, Gago-Fuentes R, Carpintero-Fernandez P, et al. Articular chondrocyte network mediated by gap junctions: role in metabolic cartilage homeostasis. Ann Rheum Dis. Jan 2015;74(1):275–84. doi:10.1136/annrheumdis-2013-204244

22. Varela-Eirin M, Varela-Vazquez A, Guitian-Caamano A, et al. Targeting of chondrocyte plasticity via connexin43 modulation attenuates cellular senescence and fosters a pro-regenerative environment in osteoarthritis. Cell Death Dis. Dec 5 2018;9(12):1166. doi:10.1038/s41419-018-1225-2

23. Varela-Eirin M, Carpintero-Fernandez P, Guitian-Caamano A, et al. Extracellular vesicles enriched in connexin 43 promote a senescent phenotype in bone and synovial cells contributing to osteoarthritis progression. Cell Death Dis. Aug 5 2022;13(8):681. doi:10.1038/s41419-022-05089-w

24. Dbouk HA, Mroue RM, El-Sabban ME, Talhouk RS. Connexins: a myriad of functions extending beyond assembly of gap junction channels. Cell Commun Signal. Mar 12 2009;7:4. doi:10.1186/1478-811X-7-4

25. Laird DW, Lampe PD. Therapeutic strategies targeting connexins. Nat Rev Drug Discov. Dec 2018;17(12):905–921. doi:10.1038/nrd.2018.138

26. Aasen T, Mesnil M, Naus CC, Lampe PD, Laird DW. Gap junctions and cancer: communicating for 50 years. Nat Rev Cancer. Dec 2016;16(12):775–788. doi:10.1038/nrc.2016.105

27. Delmar M, Laird DW, Naus CC, Nielsen MS, Verselis VK, White TW. Connexins and Disease. Cold Spring Harb Perspect Biol. Sep 04 2018;10(9)doi:10.1101/cshperspect.a029348

28. Donahue HJ, Qu RW, Genetos DC. Joint diseases: from connexins to gap junctions. Nat Rev Rheumatol. Dec 2017;14(1):42–51. doi:10.1038/nrrheum.2017.204

29. Kim JH, Ham CH, Kwon WK. Current Knowledge and Future Therapeutic Prospects in Symptomatic Intervertebral Disc Degeneration. Yonsei Med J. Mar 2022;63(3):199–210. doi:10.3349/ymj.2022.63.3.199

30. Kushchayev SV, Glushko T, Jarraya M, et al. ABCs of the degenerative spine. Insights Imaging. Apr 2018;9(2):253–274. doi:10.1007/s13244-017-0584-z

31. Totland MZ, Rasmussen NL, Knudsen LM, Leithe E. Regulation of gap junction intercellular communication by connexin ubiquitination: physiological and pathophysiological implications. Cell Mol Life Sci. Feb 2020;77(4):573–591. doi:10.1007/s00018-019-03285-0

32. Luo Y, Zheng S, Xiao W, Zhang H, Li Y. Pannexins in the musculoskeletal system: new targets for development and disease progression. Bone Res. May 6 2024;12(1):26. doi:10.1038/s41413-024-00334-8

33. Patil P, Niedernhofer LJ, Robbins PD, Lee J, Sowa G, Vo N. Cellular senescence in intervertebral disc aging and degeneration. Curr Mol Biol Rep. Dec 2018;4(4):180–190. doi:10.1007/s40610-018-0108-8

34. Aasen T, Johnstone S, Vidal-Brime L, Lynn KS, Koval M. Connexins: Synthesis, Post-Translational Modifications, and Trafficking in Health and Disease. Int J Mol Sci. Apr 26 2018;19(5)doi:10.3390/ijms19051296

35. Xu J, Shao T, Lou J, Zhang J, Xia C. Aging, cell senescence, the pathogenesis and targeted therapies of intervertebral disc degeneration. Front Pharmacol. 2023;14:1172920. doi:10.3389/fphar.2023.1172920

36. Zhao Z, Wang Y, Wang Z, Zhang F, Ding Z, Fan T. Senescence in Intervertebral Disc Degeneration: A Comprehensive Analysis Based on Bioinformatic Strategies. Immun Inflamm Dis. Nov 2024;12(11):e70072. doi:10.1002/iid3.70072

37. Gruber HE, Ma D, Hanley EN, Ingram J, Yamaguchi DT. Morphologic and molecular evidence for gap junctions and connexin 43 and 45 expression in annulus fibrosus cells from the human intervertebral disc. J Orthop Res. Sep 2001;19(5):985–9. doi:10.1016/S0736-0266(00)00072-3

38. An S, Zheng S, Cai Z, et al. Connexin43 in Musculoskeletal System: New Targets for Development and Disease Progression. Aging Dis. Dec 01 2022;13(6):1715–1732. doi:10.14336/AD.2022.0421

39. Varela-Vázquez A, Guitián-Caamaño A, Carpintero-Fernández P, et al. Cx43 enhances response to BRAF/MEK inhibitors by reducing DNA repair capacity. Nat Commun. Jul 04 2025;16(1):6168. doi:10.1038/s41467-025-60971-3

40. Ruberte LM, Natarajan RN, Andersson GB. Influence of single-level lumbar degenerative disc disease on the behavior of the adjacent segments--a finite element model study. J Biomech. Feb 9 2009;42(3):341–8. doi:10.1016/j.jbiomech.2008.11.024

