## Supplemental Table 1 for "A novel target associated with senescence and inflammatory signaling in human intervertebral disc degeneration"

**Table S1. Summary of characteristics of the donor IVD samples.**

| Patient ID | Age (Years) | Sex | Body Mass Index (BMI) | Status | Pfirrmann grade | Spinal level | Comorbidities and notes |
| --- | --- | --- | --- | --- | --- | --- | --- |
| 1 | 69 | M | 32 | DDD | 3 | L3-L4 |  |
| 2 | 69 | M | 32 | DDD | 4 | L4-L5 |  |
| 3 | 69 | M | 32 | DDD | 3 | L5-S1 |  |
| 4 | 56 | F | 39 | DDD | 5 | L4-L5 |  |
| 5 | 56 | F | 39 | DDD | 5 | L5-S1 |  |
| 6 | 7 | F | 23 | Control | 1 | T5-T6 | congenital scoli L |
| 7 | 16 | M | 18 | Control | 1 | T7-T8 | osteoblastoma T7 |
| 8 | 53 | F | 35 | DDD | 4 | L4-L5 |  |
| 9 | 71 | F | 27 | DDD in ASD | 4 | L4-L5 |  |
| 10 | 71 | F | 27 | DDD in ASD | 5 | L5-S1 |  |
| 11 | 62 | M | 33 | ADJ | 5 | L3-L4 |  |
| 12 | 62 | M | 33 | ADJ | 5 | L5-S1 |  |
| 13 | 42 | F | 19 | DDD | 4 | L5-S1 |  |
| 14 | 51 | M | 30 | DDD | 4 | L4-L5 |  |
| 15 | 60 | F | 22 | ADJ | 5 | L2-L3 |  |
| 16 | 60 | F | 22 | ADJ | 5 | L3-L4 |  |
| 17 | 43 | F | 25 | HNP | 4 | L4-L5 | in sacralization L5-S1 |
| 18 | 74 | M | 28 | HNP | 4 | L3-L4 | vertebral endplate |
| 19 | 74 | M | 28 | HNP | 4 | L3-L4 | disc tissue |
| 20 | 60 | F | 22 | ADJ | 4 | L4-L5 |  |
| 21 | 57 | F | NA | Control | 1 | T1-T2 |  |
| 22 | 42 | F | 19 | Control | 1 | C6-C7 |  |
| 23 | 53 | F | 28 | DDD | 5 | L5-S1 |  |
| 24 | 46 | M | 27 | HNP | 4 | C5-C6 |  |
| 25 | 42 | F | 29 | DDD | 5 | L4-L5 |  |
| 26 | 42 | F | 29 | DDD | 4 | L5-S1 | disc tissue |
| 27 | 42 | F | 29 | DDD | 4 | L5-S1 | vertebral endplate |
| 28 | 34 | M | 36 | HNP | 3 | L5-S1 |  |
| 29 | 59 | M | 34 | DDD | 5 | L3-S1 |  |
| 30 | 50 | F | 32 | SPL | 5 | L4-L5 | in L5-S1 sacralization |
| 31 | 71 | F | 31 | DDD in SPL | 4 | L4-L5 |  |
| 32 | 46 | M | 29 | SPL | 4 | L5-S1 |  |
| 33 | 54 | F | 29 | DDD in ASD | 5 | L2-L3 |  |
| 34 | 54 | F | 29 | DDD in ASD | 4 | L3-L4 |  |
| 35 | 54 | F | 29 | DDD in ASD | 4 | L4-L5 |  |
| 36 | 54 | F | 29 | DDD in ASD | 5 | L5-S1 |  |
| 37 | 40 | M | 29 | DDD | 5 | L5-S1 |  |
| 38 | 45 | F | 39 | DDD in SPL | 4 | L3-L4 |  |
| 39 | 42 | F | 45 | HNP | 4 | L5-S1 |  |
| 40 | 55 | F | 27 | DDD in ASD | 5 | L4-L5 |  |
| 41 | 55 | F | 27 | DDD in ASD | 5 | L5-S1 |  |
| 42 | 56 | F | 28 | HNP | 4 | L5-S1 |  |
| 43 | 55 | M | 25.39 | DDD | 5 | C5-C6 |  |
| 44 | 46 | F | 27.29 | ADJ | 3 | L4-L5 |  |
| 45 | 65 | F | 28.57 | DDD in ASD | 5 | L3-L4, L4-L5 |  |
| 46 | 42 | F | 20.93 | ADJ | 5 | C4-C5, C6-C7 |  |
| 47 | *67* | *F* | 29.17 | DDD in SPL | 4 | L4-L5 |  |
| 48 | 60 | F | 23.30 | Control | 3 | L5-S1 |  |
| 49 | 41 | M | 29.70 | DDD in ASD | 5 | L2-L3 |  |
| 50 | 46 | F | 22.10 | DDD | 5 | L4-L5 |  |
| 51 | 69 | F | 23.32 | DDD in ASD | 5 | L4-L5, L5-S1 |  |
| 52 | 33 | M | 21.22 | DDD | 4 | L4-L5 |  |
| 53 | 60 | M | 28.55 | DDD | 3 | L4-L5 |  |
| 54 | 59 | M | 27.22 | ADJ | 4 | L3-L4 |  |
| 55 | 51 | M | 27.46 | Control | 3 | C4-C5, C5-C6 |  |
| 56 | 61 | F | 29.29 | ADJ | 5 | L3-L4 |  |
| 57 | 54 | F | 32,44 | ADJ | 4 | L2-L3 |  |
| 58 | 45 | F | 30.11 | Control | 3 | L5-S1 |  |
| 59 | 54 | M | 27.76 | Control | 2 | L2-L3, L3-L4 |  |
| 60 | 70 | F | 30.29 | Control | 3 | T11-T12,T12-L1 |  |
